# Prasinovirus attack of *Ostreococcus* is furtive by day but savage by night

**DOI:** 10.1101/191478

**Authors:** Evelyne Derelle, Sheree Yau, Hervé Moreau, Nigel H. Grimsley

## Abstract

Prasinoviruses are large DNA viruses that infect diverse genera of green microalgae worldwide in aquatic ecosystems, but molecular knowledge of their life-cycles is lacking. Several complete genomes of both these viruses and their marine algal hosts are now available and have been used to show the pervasive presence of these species in microbial metagenomes. We have analysed the life-cycle of OtV5, a lytic virus, using RNA-Seq from 12 time points of healthy or infected *Ostreococcus tauri* cells over a day/night cycle in culture. In the day, viral gene transcription remained low while host nitrogen metabolism gene transcription was initially strongly repressed for two successive time points before being induced for 8 hours, but in the night viral transcription increased steeply while host nitrogen metabolism genes were repressed and many host functions that are normally reduced in the night appeared to be compensated either by genes expressed from the virus or by increased expression of a subset of 4.4 % of the host’s genes. Some host cells lysed progressively during the night, but a larger proportion lysed the following morning. Our data suggest that the life-cycles of algal viruses mirror the diurnal rhythms of their hosts.

## Introduction

The Mamiellophyceae is a class of eukaryotic unicellular green algae whose phylogenetically diverse members have been particularly successful in colonizing the world’s Oceans (1, 2). Their tiny cell sizes (3), global dispersion and ease of laboratory culture (4, 5) render them attractive as models for interdisciplinary studies in marine biology. In addition, the complete genomes of several species in the genera *Ostreococcus*, *Bathycoccus* and *Micromonas* have been characterized (6), permitting their detection in metagenomic data collected in microbial fractions of environmental seawater fractions (1). Numerous species in this group of algae are infected by prasinoviruses (1, 7), large double-stranded DNA (dsDNA) viruses in the family Phycodnavirideae. While viruses infecting *Micromonas* spp. have been known for some time (8), those infecting *Ostreococcus* and *Bathycoccus* were discovered more recently (9–13). Several of these prasinoviral genomes have now been characterized, and are typically about 200 kb long, encoding about 250 genes.

In aquatic environments in general, viruses play an important role in regulating the population of diverse phytoplankton, and affect carbon and nutrient cycling by lysing susceptible host cells(14), but much remains to be discovered about their biology. In cyanobacteria, for example, diurnal regulation of host cell lysis has been observed (15–17). Phycodnaviruses are nucleo-cytoplasmic large DNA viruses (NCLDVs) that infect many species of eukaryotic algae. The best characterized of these are PBCV-1 (*Paramecium bursaria Chlorella* virus 1), a species infecting freshwater *Chlorella*, that are also symbionts of the ciliate *Paramecium bursaria,* (18) and *Emiliania huxley* viruses(19) that infect the marine haptophyte unicellular alga *Emiliana huxleyi*, well-known for its extensive oceanic blooms.

The life-cycle of *Ostreococcus tauri* virus 5 (OtV5), with its typical icosahedral particle morphology, 8-hour long latent period and small burst size of 25, cultured with its host in continuous light, was first described by Derelle *et al* (9). Numerous studies describing viral growth in related prasinoviruses infecting *Micromonas* have been made previously using a day-night cycle (20–27), and recently Demory *et al* (28) revealed temperature to be a key factor in these interactions, but detailed molecular analyses were not the main objective of these studies. In the present study, we aimed to re-examine growth of OtV5 in a more natural light regime (12h light / 12h dark) and to characterize gene expression from the host and algal genomes by RNA-Seq transcriptomic analyses.

## Results

### Growth of host cells and virus after infection at different times

A partial synchronization of *O. tauri* growth was previously reported under a 12:12 light/dark (L:D) cycle (29). In these cultures, cells were in G1 phase during most of the light phase and progressively entered in S phase and mitosis at the end of the day. The division of the population occurred during a period of 2 hours before and 2 hours after the light to dark transition. In such synchronized cultures, the effect of inoculating cultures at different times during the day was thus tested in a preliminary experiment, to find the best time to inoculate the cultures. For that purpose, *O. tauri* cultures were infected with OtV5 every 2 hours during the light phase (Fig. 1). Cell lysis was almost complete at the end of the second day (36 hours later) when the infection occurred in the first 4 hours of the light period. In contrast, when infection occurred later, for example after 10-12 hours of light, no cell lysis was observed at 36 hours after the start of the experiment and lysis was delayed until the next day.

**Fig 1.**
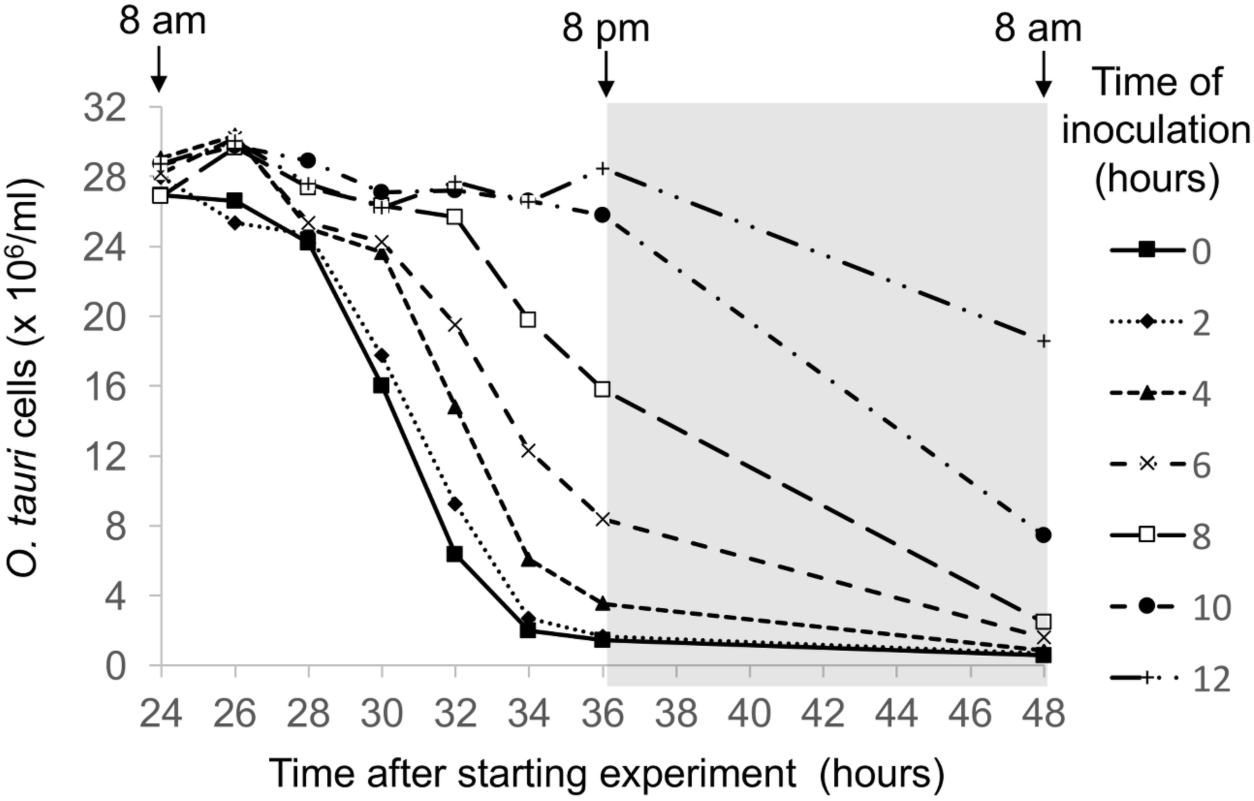
Time course of lysis of *O. tauri* cultures partially synchronized by a 12:12 light:dark cycle and inoculated with OtV5 (MOI 5) at different times during the previous day, as indicated in the adjacent key. Note that almost complete lysis of cells occurred at 36 hours post inoculation (hpi) only when cultures were inoculated on the previous day at 8 am (time zero, filled squares with continuous line) or 10 am (time 2, fine dotted line with filled diamonds).

### Virus-host infection dynamics

In order to have a complete viral lysis cycle within two working days, we infected cells using a multiplicity of infection (MOI) of 10 per cell one hour after “dawn”, giving 11 hours of light before the dark period. In these conditions, infected cells did not start to lyse at 8 hours post inoculation (hpi) as observed previously under continuous light (9), but remained intact until after “nightfall”. Uninfected and infected cells started to divide at 7 hpi, but cell growth in infected cultures was strongly reduced after 9 hpi (Fig. 2). Control cells divided 7–17 hpi, doubling in cell density, but only about 40% of infected cells divided, reaching a maximum at 13 hpi, then declining slowly until 1 hour after “dawn”, when about 80% of the remaining cells lysed in the period 23–25 hpi (Supplemental Fig. S1A). By 27 hpi, the “infected” population was reduced to about 7% of the control.

**Fig 2.**
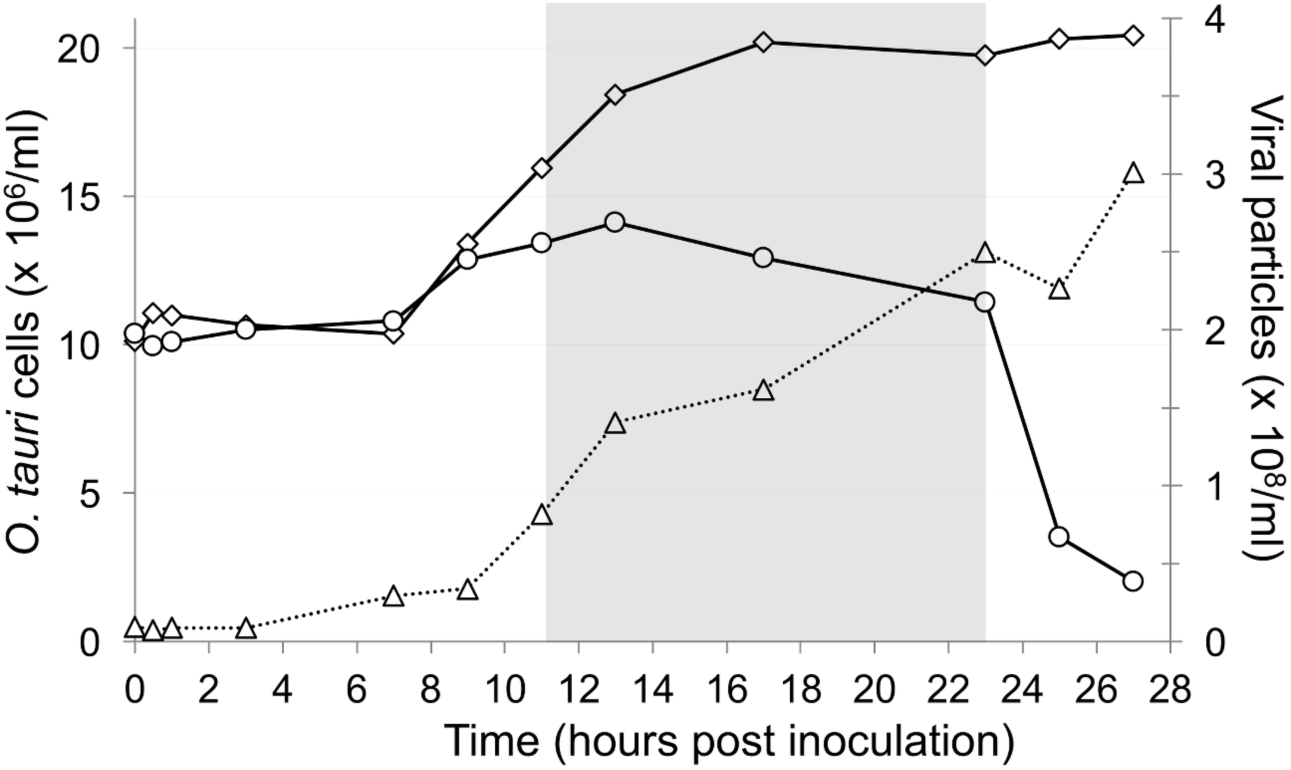
Growth curves of *O. tauri* cultures and OtV5. Open diamonds: uninfected *O. tauri*, open circles: *O. tauri* infected by OtV5 and open triangles: OtV5 production.

In inoculated cultures, given that the MOI was 10, we observed that most viruses adsorbed to each cell immediately after inoculation, since the density of particles measured by flow cytometry appears to drop to a tenth of that in the inoculum. Few viruses were released from host cells by 9 hpi, but many more were released in the period 9–13 hpi, and the viral population continued to increase until the end of the experiment, when the total number of virus particles was 25–30× higher than the number of host cells inoculated at 0 hpi, in good agreement with the burst size of 25 calculated previously (9).

## Differentially expressed host genes

mRNAs of all samples were analysed using RNA-Seq technology and host and viral gene expression was compared at each time point between control and infected cells. With the parameters used in the analysis (see Methods), only 323 host genes were significantly differentially regulated at any one time point using the chosen analytical parameters (Table 1, “Methods” and Supplemental Table S1).

**Table 1.**
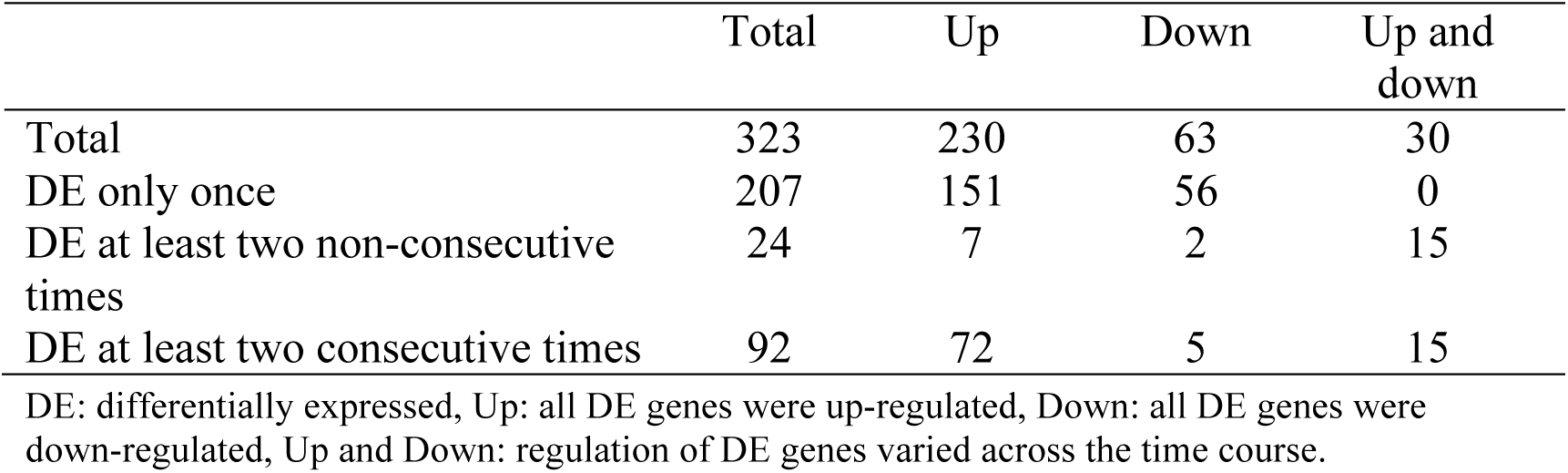
Numbers of differentially expressed *O. tauri* genes.

|  | Total | Up | Down | Up and down |
| --- | --- | --- | --- | --- |
| Total | 323 | 230 | 63 | 30 |
| DE only once | 207 | 151 | 56 | 0 |
| DE at least two non-consecutive times | 24 | 7 | 2 | 15 |
| DE at least two consecutive times | 92 | 72 | 5 | 15 |

DE: differentially expressed, Up: all DE genes were up-regulated, Down: all DE genes were down-regulated, Up and Down: regulation of DE genes varied across the time course.

**Supplemental Table S1** Host genes differentially expressed once or more between control and infected cultures at different times after inoculation. Log2-fold changes are shown only if the number of raw reads exceeds 100 in both control and infected treatments.

Given the high number of sampled time points in this experiment (24 mRNA libraries were sequenced), no replicates were done. To palliate this, only genes whose expression was differentially regulated at two or more consecutive sampling times were retained. Application of this criterion decreased to 92 the number of host genes which were considered to be regulated (Fig. 3 and Table 1).

**Fig 3.**
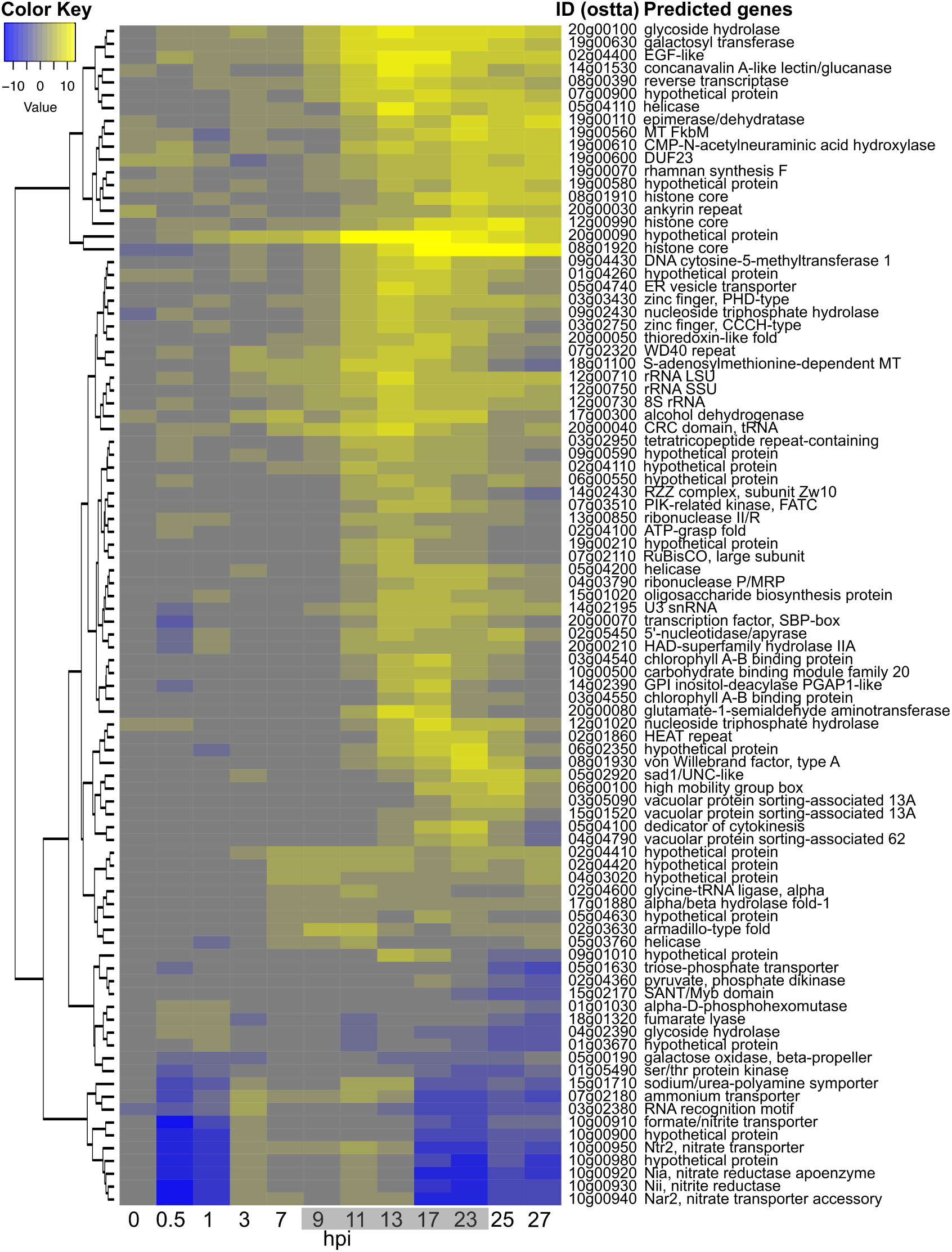
Differentially transcribed *O. tauri* genes during a 27-hour infection time course. Time (hpi) is shown along the abscissa, with time points sampled in the dark shown with a grey background, and rows represent DE genes clustered according to log2-fold changes in expression (Color Key)(see Supplementary Table S2 for a detailed list of genes).

**Supplemental Table S2** Host genes differentially expressed at least 2 consecutive time points between control and infected cultures. Log2-fold ratios and FPKM values are shown for all time points.

*See Excel spreadsheet*

Most of them (72) were only up-regulated whereas 5 were only down-regulated. Fifteen other genes were also regulated in the opposite direction at least once in the course of the experiment, albeit 13 having a consecutive regulation at two successive times in the same orientation (up-or down-) (Table 1). Most of the regulated genes were individually dispersed in the genome except for a cluster of 7 genes, including the nitrate/nitrite transporter/reductase cluster previously described (30), and a group of genes on chromosome 19 (see below). Among the 92 genes, 77 (80%) had a potential identified function (Supplemental Table S2), but no clear metabolic pathways could be identified except for the nitrate/nitrite cluster mentioned above and present on chromosome 10 (Supplemental Fig. S2). This tandem organisation indicated a possible selective pressure for optimization of nitrate uptake and assimilation by *O. tauri*, although experimental evidence for such a coordinated expression of these genes is currently lacking.

**Supplemental Fig. S2**. Distribution of *O tauri* differentially transcribed genes across the genome. Numbers underneath chromosome numbers indicate the number of genes encoded by that chromosome. Large black rectangles represent chromosomes, each rectangle being a composite view of the 12 time points during the OtV5 infection stacked from bottom to top. On chromosome 10, for example, the nitrogen simulation cluster of genes (blue arrowhead) can be seen as under-expressed (green), then over expressed (red), then under expressed in the final stages, reading from bottom to top of the rectangle.

Interestingly, in our experiment their expression was first strongly inhibited during at least the first hour post infection, then, up-regulated up to the same level of the control until 17 hpi, and, finally, again strongly inhibited (Fig. 3).

Ribosomal RNA gene transcripts were much more abundant in infected cells than in healthy cells during the infection process (Fig. 3).

## Early fluctuations in host transcript abundance

Disregarding the above requirement for DE in the same direction at two consecutive points, we observed that a small number of host genes showed up-down or down-up regulation at two early time points after infection. Eight genes showing DE at one time point were up-regulated at 0-1 hpi then down regulated (up-down) and 7 genes showed down regulation at 0 to 1 hpi then up regulation (data not shown). Most of these proteins are predicted to have regulatory functions (4 transporters, 3 nucleic-acid binding, 2 transcription factors, 2 unknowns, 1 kinase, 1 ATPase).

## Expression of viral genes

All of the viral genes were expressed during the life-cycle, except for most of those present in the long terminal inverted repeats (TIRs). Clustering the data (Fig. 4) revealed successive functional groups of genes. The expression pattern of viral genes in infected cultures occurred in two phases (Fig. 4).

**Fig 4.**
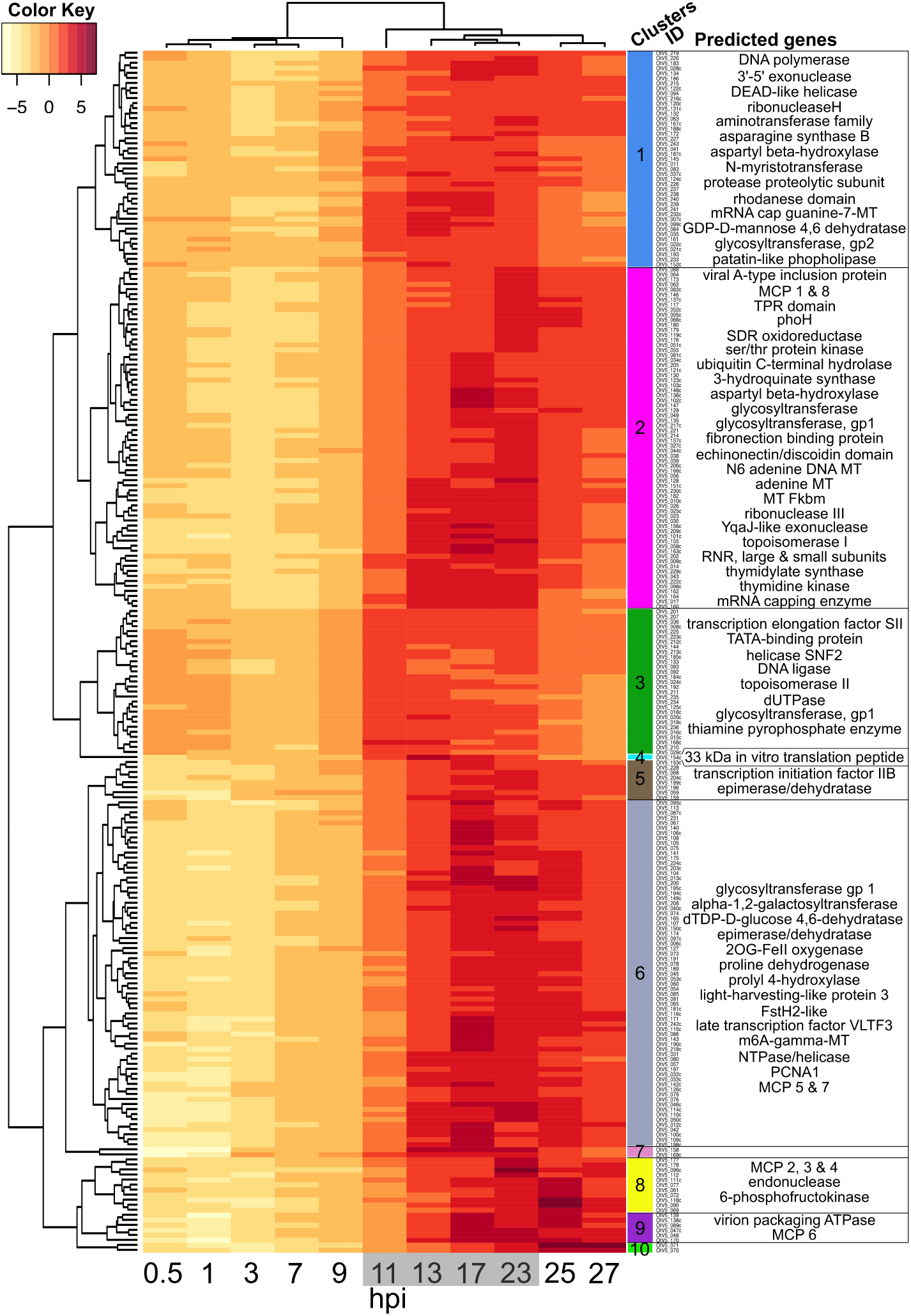
Timing of OtV5 relative gene transcription during infection. Time (hpi) is shown along the abscissa, with time points sampled in the dark shown with a grey background. Rows represent OtV5 genes clustered according to the variation in their relative expression the over time (left dendrogram) and by the relative expression pattern of each sample (above dendrogram). The colour key shows regularised log (rlog) transformed gene fragment counts centred to the mean of each gene (row means). (see "Methods" and Supplemental Table S3 for a detailed list of genes).

**Supplemental Table S3** Expression of OtV5 genes at different time points during the infection, grouped according their expression profiles (see also Fig. 4 and Methods). The genes in TIRs (4 genes at each end of the genome) showed no or negligible expression and were not included.

*See Excel spreadsheet*

Phase I from 0–9 hpi with low viral transcription (<6% of reads mapped to OtV5)corresponding to the start of cell division before the light/dark transition and phase II occurring after from 11–27 hpi with high viral transcription (up to 66% of reads mapped to OtV5). During the first phase in the light, two clusters of phase I genes (clusters 3 and 1) stood out as more strongly expressed than others. Cluster 3 was strongest, concerning genes mainly involved in controlling transcription initiation and nucleotide processing, whereas cluster 1 contained a mixture of functions. Both of these clusters contained genes involved in DNA replication. The majority of phase II viral gene expression can be seen to occur probably when host DNA replication has been completed(29) at 13–23 hpi, in clusters 2 and 6, whereas clusters 8, 9 and 10 contain genes that are highly expressed very late in infection. Phase II also contains genes classically involved in late virus particle development, such as major capsid protein-like (MCP), viral A inclusion body protein and virion packaging ATPase. At the end of phase II, all viral genes were expressed, except for 3 of the 4 genes in each TIR (Supplemental Figure S3).

**Supplemental Fig. S3** The distribution of transcript abundance (FPKM) of OtV5 genes during the time course of infection (hpi shown on right). The OtV5 genome is represented from left to right, one gene per column on the heat map. Six out of 8 of the genes in the terminal inverted repeats (TIR) were not expressed, so that blue columns appear at genome extremities.

The most expressed viral gene was annotated as a 33 kDa *in vitro* peptide translation (Supplemental Table S3) whose function is not clear but which was also been reported to be massively expressed in the closely related Chlorella viruses(31). This gene was highly expressed throughout the infection. Among the 8 major capsid protein (MCP)-like genes, copies 1–7 began to be expressed late in the second phase, between 23 and 25 hpi, whereas copy 8 was expressed earlier (Fig. 4, Supplemental Table S3).

Several genes with similar predicted functions are present in both the virus and the host genomes. The regulation of their respective expressions showed two patterns. The first pattern showed highest expression of the host gene during the light phase and highest viral gene expression in the dark, when the host gene counterpart expression was low. The virus thus appeared to be autonomous for some of the functions necessary for its growth in the night (Fig. 5). For example, this was observed for the two subunits of the ribonucleotide reductase and for the DNA polymerase (Fig. 5). In the second pattern, the expression of the host genes was inhibited in the infected cells compared to the control whereas the virus genes were expressed (Fig. 5,), again appearing as a compensation of the host gene inhibition. This is illustrated by the asparagine synthase gene (Fig. 5) but was observed for several other genes such as the proline dehydrogenase and the acetolactate synthase.

**Fig 5.**
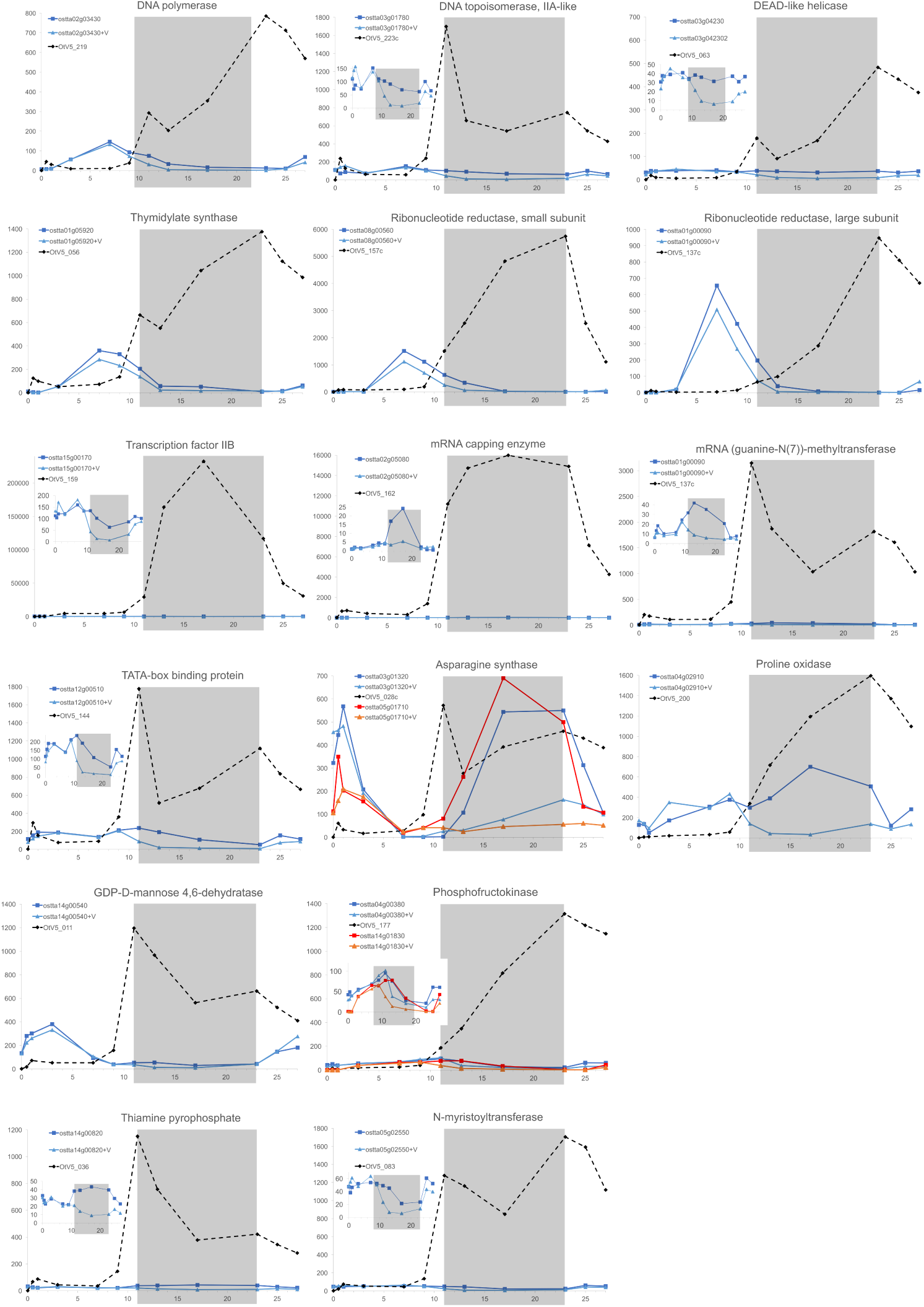
Expression (ordinates: FPKM) of healthy (square data points, darker colours), infected (triangular data points, lighter colours) *O. tauri* and OtV5 (diamonds, dashed lines) genes sharing a similar putative function. Abscissae: time (hours post-inoculation). Inoculation with the virus was done at t = 0, one hour after daytime (light) started. The grey shaded area indicates the night (dark) period. Genes are identified by their accession numbers in public databases (top left in each panel, +V: virus-inoculated cultures).

## Discussion

Several previous detailed reports on the life-cycles of large DNA viruses infecting green microalgae have been done in continuous illumination (9, 32–36), which promotes rapid growth of the host and virus, but since all of these algae have evolved in a diurnal cycle, we decided to perform this study in a 12h light and 12h dark “day” and “night” cycle. In healthy cells, under these conditions, the general pattern of gene transcription is quite different in the daytime, when photosynthesis is in progress, and in the night, when stored energy is being used, this rhythm being observed both in the laboratory (37) and in the environment (38).

Whereas under continuous illumination there was a burst of viruses released at 8 hpi (9), in the light-dark cycle, the timing of the host cell lysis was variable, with some cells lysing during the night, but most of the cells dying after illumination of the cells the following morning (Fig. 2). Under these conditions, it does not really make sense to think of the “burst” time as a fixed period. It probably also varies according to the temperature and in nature, the seasons in temperate latitudes. Several authors have investigated the effects of host cell cycle (39) or different environmental variables on viral life-cycles (20, 40–43), but appropriate tools were not available or not used for molecular analyses in these species. Using gene-specific probes or biochemical analyses, *E. huxleyi* viruses were shown to affect certain host metabolic pathways(44, 45), but diurnal variations were not discernible in this system. Our observations agree well with those of Brown et al (22) on the related prasinovirus *Micromonas* virus MpV-Sp1, who observed a peak of viral production about 24 h after infection. Furthermore, these authors showed that host cell lysis was delayed in prolonged dark periods and confirmed their observations on host cell densities and virus production using molecular probes.

In natural populations of phytoplankton as in culture, cell growth responds strongly to light/dark periodicity (38, 46). Our data support the notion that viral gene transcription is rather quiescent during the day and increases rapidly at the onset of the dark when host DNA replication is being completed, thereafter remaining strongly expressed. Many of the genes that were significantly expressed in the quiescent phase are abundant in the active later phases, suggesting that their quiescent expression may reflect to some extent leaky general suppression levels. However, the heat map clustering revealed that viral genes for nucleic acid processing and transcription do appear to be more abundant than other messages in the first phase (Fig. 4, clusters 1 and 3), although these genes continue to be expressed among the late genes. For example, transcription factor IIB (TFIIB), a conserved gene in eukaryotes and many large DNA viruses that is part of the core transcriptional machinery (47) was very highly expressed in the night (Fig. 5). In the night, viral genes probably essential for the late stages of viral growth appeared to compensate for gene functions that were normally turned down in the night, including functions probably important for DNA replication and amino acid metabolism, while transcripts likely encoding virion assembly and glycosylation were highest in the latest time points (Figs. 4 and 5). Although only arginine synthase and proline oxidase showed significantly different levels between control and infected at one time point (Supplemental Table S1), insufficient for our requirement of consecutive times, the strength the coordinate swings in expression shown in Fig. 5 for 16 genes clearly intimates that viral metabolism predominates, justifying our approach of numerous sampling times. Some host amino acid synthesis genes normally expressed in the dark were turned down in the dark in virus-infected cultures, but their viral counterparts were then up-regulated (Fig. 5). Viral proline oxidase was probably acquired from its host genome (9), is known to produce ATP during stress responses in eukaryotes (48, 49), and is a possible source of energy for the virus. Phosphofructokinase is a key enzyme controlling the production of energy through glycolysis(50) and viral transcript levels in the night rose to over an order of magnitude higher than those of the host (Fig. 5).

## Ribosomal RNA overexpression

Although our extraction procedure was designed to isolate polyadenylated messenger RNA, some ribosomal RNA genes (rRNA), which are always abundant in RNA extractions of active cells, were represented in our data. There is increasing evidence that rRNA transcripts can be polyadenylated in eukaryotes, including algae in the same phylogenetic order as *Ostreococcus*, such as *Micromonas* (51). rRNAs were over-represented late in infection in *O. tauri*, compared with the control. At least three explanations are possible for this. Firstly, it may result because the ribosome is a large and relatively stable subcellular structure that might persist better than the other cytoplasmic RNAs during the late viral infection thereby preferentially protecting ribosomal RNAs that lie buried within it. Much of the available cellular RNA pool is likely to be used by the viral ribonucleoside-diphosphate reductase, an enzyme with two subunits that all prasinoviruses encode, to permit synthesis of prasinovirus DNA. This viral enzyme continued to be highly expressed during the night (Fig. 5), when the equivalent host genes were shut down. Secondly, an over-expression of ribosomal RNA may be induced by the virus. U3, an RNA probably transcribed by RNA polymerase III (52) and essential for the first step of pre-rRNA processing (53) is apparently overexpressed during late viral infection. *O. tauri* RNA pol III is normally constitutively expressed as it required for many basic cellular functions (54), and indeed there is no significant difference observed in the expression of its controlling repressor, ostta05g03220, that encodes the orthologue of Maf1(55). The apparent abundance of U3 suggests that it may not be dislodged from the ribosomal RNA for processing. The proteins UTP14 and DHR1 are required to dislodge U3, (56), but in *O. tauri* the putative orthologues of these genes (ostta04g00770 and ostta05g03760, respectively) are not induced. Only 3 of the 161 annotated *O. tauri* ribosomal proteins were modestly over-expressed at one time point, the other being under-expressed (Supplemental Table S4). This would thus result in overproduction of unprocessed host ribosomal RNA precursors, potentially providing OtV5 with a rich source of nucleic acids by their degradation. Thirdly, in yeast, where the dynamic, energetically demanding and complex process of ribosome biogenesis has been studied in detail (57), nutrient starvation or stress are known to shut down the synthesis of ribosomes *via* the conserved global regulatory TOR (target of rapamycin) pathway at the stage of initiation of transcription or pre-rRNA (57, 58). However, in our system rRNA appears to accumulate and its processing occurs in an apparently normal way up to 23 hpi (Supplemental Fig. S1). Recently Kos-Braun et al (59), demonstrated an alternative pathway for blockage of rRNA processing at a later stage at during the diauxic growth phase in yeast. When glucose is no longer available casein kinase 2 (CK2, an orthologue of ostta12g02550 in *O. tauri* (60)) can phosphorylate TOR1, and partly processed rRNA products can accumulate in a resting (G1 or G0) stage. While this type of control also leads to accumulation of rRNA, Kos-Braun et al show that the 5S moiety in yeast remains attached to the large rRNA subunit precursor, whereas in *O. tauri* the accumulated rRNA look normal.

**Supplemental Table S4** Transcript abundancies (FPKM) for all host genes at different times in healthy control cultures (light blue background) and cultures inoculated with OtV5 at time zero (light red background)

While our data favour the second hypothesis, further work is required to study this process in more detail, since it may be a pivotal switch governing acquisition of sufficient cellular metabolites to resource the biosynthesis of large viral genomes before the host cell bursts. A least two of the control steps of host rRNA production might occur by protein phosphorylation (phosphorylation of TOR by its controlling proteins either at the stage of pre-RNA initiation or at a later stage (59)), and were out of the scope of the current study. More precise analysis of processing at the 5’ part of the pre-rRNA (the position of U3 binding) would also be desirable.

## Nitrogen assimilation

The uptake and conversion of nitrate to its reduced form required for synthesis of amino acids is a complex and energetically demanding process (61). The expression of genes involved in the assimilation of nitrate, the only source of nitrogen in our culture medium, and many others of the N assimilation pathway, are strongly differentially expressed throughout the course of infection, being firstly repressed, then induced and finally repressed (Fig. 3 and Supplemental Fig. S2). These include numerous genes clustered together on chromosome 10 and a few genes scattered on other chromosomes. This is striking, because it is not related to the nitrogen sources available in the medium. In addition, the none of the 3 cyclin-dependent protein kinase genes shown to be involved N assimilation responses(62) showed differential expression, suggesting that this response is not functioning. There is an adequate level of nitrate in the culture medium used, (no ammonium provided in L1, see “Methods”), so nitrate uptake and nitrate reductase genes should be highly active, as they are in the control. There are several possible non-mutually exclusive reasons for this repression, which might either be initiated as a host defence response or be the result of a virally encoded products influencing N assimilation by this pathway.

Reduction of nitrate via nitrate reductase also leads to production of nitric oxide (NO)(63, 64), a signalling reactive oxygen species (ROS) active in diverse species (65, 66), including algae (61) that is known to heighten the cellular defence responses of cells to stress (66–68). It is required for resistance to viruses in of *Arabidopsis* (69) and rice (70) and ROS are also known to modulate the response of *E. huxleyi* to viruses(45). Since the Nitrogen and Carbon / Phosphorous ratio for small green algal structural and metabolic requirements far exceeds that of the nucleic-acid rich large DNA viruses (71) and the cell is doomed to lyse, it may be advantageous for the virus to divert the resources usually used for protein synthesis towards nucleic acid synthesis, at the same time lowering the chance of detection by NO signalling that would initiate host defences. If the TOR complex is targeted by the virus as suggested above, and as shown recently in other host-pathogen systems (72, 73) this might also lead to TOR-controlled repression of the nitrogen assimilation genes (74). The coordinated regulation that we observed suggests the involvement of a global regulator, with opposing forces governing this control, provoking a strongly fluctuating response. However, the recent demonstration that certain prasinoviruses have acquired host genes that permit uptake of reduced nitrogen(75) suggests that this resource may also be limiting during infection, and favours the notion that suppression of NO signalling is the reason for decreasing the uptake of nitrate.

## Is the replicative form of OtV5 chromatinized?

In several other host-virus systems, chromatinization of viral DNA that enters the nucleus is known to occur rapidly once the viral DNA enters the nucleus (76–78). The replicative form of OtV5 has not yet been investigated, but it very likely has a nuclear phase during its infection cycle as OtV5 lacks a DNA-dependent RNA polymerase to transcribe viral genes (9). Herpes Simplex Virus (HSV) for example, is a dsDNA virus that probably replicates in the nucleus and is packaged in capsids as a linear molecule in the cytoplasm. During the HSV lytic cycle the viral genome circularizes and nucleosomes form along its genome (79) in a highly dynamic way that is modulated by a viral transcription factor (80). The strong induction of all host histone core genes observed throughout the OtV5 life-cycle strongly suggests that the OtV5 genome is chromatinized during replication of the viral genome, and that viral replication continues throughout the dark cycle, when many photosynthesis-dependent host processes are shut down (81). Host S-adenosylmethyltransferase, an enzyme required for the majority of processes that modify DNA, RNA, histones and other proteins, including those affecting replication, transcription and translation, mismatch repair, chromatin modelling, epigenetic modifications and imprinting (82), was overexpressed in a similar way, suggesting that any of these pathways might be induced during viral infection. Its continued expression, also during the night, is likely necessary for the numerous pathways required for virus production.

## Induction of reverse transcriptase

The *O. tauri* reverse transcriptase gene ostta08g00390 was strongly induced (over 4 consecutive time points, and up to 420-fold at 13 h post-inoculation, (Fig. 3, Supplemental Table S2), during the period when cell division is expected to occur (at the end of the day, from 2 h before dark then for the following 6 h). This gene is the predicted replicase/integrase of a putatively complete type I transposon (30, 83, 84) that is not usually active in healthy *O. tauri* cells. At 7-13 hpi we observed a strong increase in the transcription of this gene. We hypothesize that the increase in transcription of this reverse transcriptase may be activated by the cellular stress response caused by OtV5 attack, that may in turn activate transposition itself and the repeat retrotransposon in miniature (TRIM) on chromosome 19, leading to chromosomal rearrangements and possibly to activation of certain genes on chromosome 19 whose expression continues late in infection in those cells that subsequently become resistant to viral attack. Yau et al (2016)(84) observed rearrangements on chromosome 19 and overexpression of genes on this chromosome in cell lines that had become resistant to OtV5 infection, and the karyotypes of these strains also suggest possible rearrangements and/or translocations on chromosomes 19. This may additionally explain the presence of DNA very high in the PFGE gel since long reads of that chromosome by reverse transcriptase from transposon LTR might generate DNA intermediates that would not enter the gel (85–87). Recently, Blanc-Mathieu et al(88) revealed astonishing variability in the structure of chromosome 19 in natural populations of *O. tauri*. Whether or not rearrangements of this chromosome contribute to the acquisition of viral resistance is not yet clear, and will be a subject for future investigations.

## Host genes induced very late

While most of the host and viral differentially expressed genes showed increased transcription just after the beginning of the night time (Fig. 3), a time when we expect host DNA replication to be underway, surprisingly, a few host genes showed a second period of induction very late in infection, during the 2^nd^ half of the night and morning of the next day, 17–27 hpi. Since several of them were also observed to be induced in OtV5-resistant lines of *O. tauri* (84) we compared the host genes identified in both experiments as being differentially expressed. Twenty*-*six genes were found to be differentially expressed at some stage in both of the analyses, and the expression of 11 of them were strongly up-regulated in the last 13–27 hpi of the experiment. Since the majority of these genes (6/11) were located on the viral immunity chromosome first described by Yau et al (84), we hypothesize that this expression originates from a sub-population of resistant cells that have differentiated from the bulk of the susceptible cells, the latter being condemned to lyse and release viral progeny.

In summary, we have shown that in a natural light regime the life-cycle of prasinoviruses in *Ostreococcus* in culture is biphasic, remaining quiescent by day but becoming full-scale at night, when new virus particles arise steadily at first and then rather suddenly in the morning. During the night 239/247 (96.8%) of predicted viral genes are transcribed, and 323/7749 (4.2%) host genes are differentially expressed at some stage, the great majority (71%) being up-regulated, in response to the viral attack. However, the pattern of host gene expression in the final phase of infection already suggests that a small population of host cells were adapting to become founders for resistance to OtV5. Detailed knowledge of host-virus interactions will be necessary for advancing our understanding of the everlasting war between hosts and their viruses in aquatic environments.

## Methods

### Culture conditions and growth measurements

The host strain *Ostreococcus tauri* RCC4221 (30, 83, 89) and the prasinovirus OtV5 (9) were used in all experiments. Cultures were grown in L1 medium (Bigelow Laboratory, NCMA, USA) diluted in 0.22 μm filtered seawater under a 12/12 light/dark cycle (100 μmole photon/m^2^s^-1^). Cell and viral counts were performed on a FACScan flow cytometer (Becton Dickinson, San Jose, CA, USA). *O. tauri* cells were counted according to their right-angle scatter and their red fluorescence emission due to the chlorophyll A pigment (90). OtV5 counts were determined by their right-angle scatter and their fluorescence after SYBR green I staining (91). For preparation of large quantities of viruses, five litres of an *O. tauri* exponentially growing culture (approx. 5.10^7^ cells.ml^-1^) was inoculated with an OtV5 lysate. Lysed cultures were centrifuged at 8000 *g* for 20 min at 20°C and then passed sequentially through a 0.22 μm filters to remove large cellular debris. Virus filtrates were concentrated by ultrafiltration with a 50K MW size cut-off unit (Vivaspin 15 Turbo, Sartorius) to a final volume of 5 ml. Concentration of infectious particles was determined by a serial dilution assay.

To test for the effect of viral infection at different times during the day, *O. tauri* cultures in exponential growth phase were infected with purified OtV5 at a multiplicity of infection (MOI) of 5 and cell counts measured over 48 h.

To perform the differential expression analysis, an *O. tauri* culture was acclimatised such that cell density doubled every day from 10^7^ to 2.10^7^ cells/mL by flow cytometer counting and diluting the culture daily for 10 days. After this period of acclimation to maintain cultures in this rhythm of growth, 1.5 L of *O. tauri* culture was prepared, counted by flow cytometry, adjusted to a cell concentration of 10^7^ cells ml^-1^ by addition of L1 medium and half of the culture was infected with OtV5 one hour after the beginning of the light phase with OtV5 at a MOI of 10. The cultures were then split into control and infected, comprising 12 × 100 mL flasks for each condition. At 12 different times between 0 and 27 hours post inoculation (hpi), control and infected flasks were sampled to measure cell and viral densities by flow cytometry and cells harvested for RNA extraction (Fig. 2 and S1).

### RNA extraction and sequencing

For RNA extraction, 50ml of cells were harvested by centrifugation at 8,000 *g* for 20 min at 20°C. The pellets were then flash frozen in liquid nitrogen and stored at -80°C. Total RNA was extracted using the Direct-zol™ RNA kit (Zymo Research), and checked for quality (Supplemental Table S1B). Selection for polyadenylated RNA, library preparation and sequencing was performed commercially (GATC Biotech AG., Konstanz, Germany). RNA libraries were sequenced on the Illumina Hi-Seq 2000 platform by multiplexing all samples on a single flowcell lane, which generating paired end reads of 101 bp in length. RNA sequence reads were checked for quality using FastQC.

### Differential gene transcription analysis

Transcriptome read pairs (fragments) were aligned using TopHat2 (92)(alignment parameters: -i 17 -I 3500, -G) to the annotated genome sequence of *O. tauri* RCC4221(83) and OtV5 (9). The counts of fragments aligning to each gene were determined using the htseq-count function of HTSeq (93) with parameters (-m intersection-nonempty). Fragments per kilobase of exon per million reads mapped (FPKM) were calculated for visualization of the expression of individual genes of interest and of OtV5. Differential gene expression analysis and data visualisations were performed in the R statistical environment (https://www.r-project.org/). Differential host gene expression analyses were performed on fragment count tables using the R package DESeq(93) to detect genes involved in OtV5 infection. Host gene transcription from each sampling time point was compared between control uninfected and infected cells using the DESeq function accepting genes as significantly differentially transcribed with adjusted p-value < 0.1. Candidate host genes involved in viral infection were accepted if > 100 reads were assigned to the gene and if they were differentially transcribed in at least two consecutive time points. Heatmaps and hierarchical clustering (euclidean distance) were produced using heatmap.2 on the log2-fold change values of *O. tauri* DE genes and the row means centred regularised log (rlog) (from DESeq2 function) transformed fragment counts of OtV5 genes with non-zero fragment counts. OtV5 clusters of genes that covaried in their relative expression over time were designated using the cuttree function at h=5.

## Accession numbers

*O. tauri* RCC4221 chromosome sequences can be found under the GenBank accession numbers CAID01000001.2 to CAID01000020.2 and gene annotations are also available from the Online Resource for Community Annotation of Eukaryotes (ORCAE) under *Ostreococcus tauri* V2. The updated genome sequence and annotation of OtV5 is available under GenBank accession EU304328.2. Transcriptomic data used in this study is available under project accession PRJNA400530.

## Acknowledgments

We thank D. Pecqueur and S. Salmeron for use of the of the cytometry platform, Céline Noirot (Genotoul platform, Toulouse) for advice on data analysis, and the GENOPHY team (Banyuls sur Mer) for discussions. We are grateful for financial support from the French National Resaerch Agency (grants REVIREC ANR-12-BSV7-0006-01 and DECOVIR ANR-12-BSV7-0009).

## Author contributions

HM & ED planned the experiments, ED did experimental work, ED and SY did bioinformatic analyses, all authors were involved in interpreting the results, NG, SY and HM wrote the article.

## Additional information

Supplemental information is available online. Correspondence and requests for materials should be addressed to H.M. or N.H.G.

---

**Supplemental Fig. S1** A: Visualisation of *O. tauri* cells by flow cytometry in healthy (control) or infected (inoculated at 0 hpi) from samples taken at different times during host and viral growth during the RNA-Seq analysis. Healthy *O. tauri* cells are seen as red fluorescent points clustering in the window shown, whereas lysing cells can be seen as dark points underneath the window with reduced fluorescence and side scatter that begin to appear in infected cultures at 9 hpi, becoming suddenly stronger at 25 hpi and rising to maximum at 27 hpi.

B: Electropherograms of RNA extractions from in healthy (control) or infected (inoculated at 0 hpi) cultures on an Agilent 2100 Bioanalyzer, dark bands showing mainly abundant ribosomal RNAs. In infected cultures the extracted RNA is partly degraded after 23-27 hpi, when most of the host cells are lysing. M – molecular weight marker track.

